# Effects of photobiomodulation on the redox state of healthy and cancer cells

**DOI:** 10.1101/2020.09.07.285908

**Authors:** Clara Maria Gonçalves de Faria, Heloisa Ciol, Vanderlei Salvador Bagnato, Sebastião Pratavieira

**Affiliations:** São Carlos Institute of Physics - University of São Paulo, São Carlos, SP, Brazil; Faculty Fellow at the Hagler Institute for Advanced Study and Visiting Professor at the Department of Biomedical Engineering - Texas A&M University, College Station Texas - USA

**Keywords:** photobiomodulation, optical redox ratio, mitochondria, NADH, FAD

## Abstract

Photobiomodulation (PBM) uses light to stimulate cells. The molecular basis of the effects of PBM is being unveiled, but it is stated that the cytochrome-c oxidase enzyme in mitochondria, a photon acceptor of PBM, contributes to an increase in ATP production and modulates the reduction and oxidation of electron carriers NADH and FAD. As it can stimulate cells, PBM is not used on tumors. Thus, it is interesting to investigate if its effects correlate to mitochondrial metabolism and if so, how it could be linked to the optical redox ratio (ORR). To that end, fibroblasts and oral cancer cells were irradiated with a light source of 780 nm and a total dose of 5 J/cm^2^, and imaged by optical microscopy. PBM down-regulated the SCC-25 ORR by 10%. Furthermore, PBM led to an increase in ROS and ATP production in cancer cells after 4 h, while fibroblasts only had a modest ATP increase 6 h after irradiation. Cell lines did not show distinct cell cycle profiles, as both had an increase in G2/M cells. This study indicates that PBM shifts the redox state of oral cancer cells towards glycolysis and affects normal and tumor cells through distinct pathways. To our knowledge, this is the first study that investigated the effects of PBM on mitochondrial metabolism from the initiation of the cascade to DNA replication. This is an essential step in the investigation of the mechanism of action of PBM in an effort to avoid misinterpretations of a variety of combined protocols.

## Introduction

Photobiomodulation (PBM) has been used for decades for wound healing, tissue regeneration, analgesia, inflammation reduction, osteoarthritis, reducing edema on lymph nodes, and muscle relaxation, among others (1, 2). However, it is a developing field which results in partial acceptance and recognition from authorities in biomedical science, professionals and scholarly journals (3). It encompasses a variety of reactions caused by non-ionizing and non-thermal light absorption in tissues and cells, resulting in a physiological response according to tissue stimulation. However, its effects are still unclear, particularly on premalignant and malignant cells. One of PBM most popular applications, due to its effectiveness, is the prevention and management of oral mucositis in head and neck squamous cell carcinoma (HNSCC) patients (4, 5). Still, a recent systematic review, including 13 papers, demonstrated that the data does not support a definite conclusion of PBM impact on HNSCC cells, despite many studies on the topic (4). Among the challenges are the wide variety of study designs, PBM protocols and the limited type of assays performed, where cell proliferation and viability are the primary ones.

Evidence indicates that the PBM cascade of events begins with cytochrome c oxidase (COX), the fourth protein complex in the mitochondrial electron transport chain and primary photoreceptor of red and near-infrared light (6–8). The energy absorbed by COX changes the mitochondrial potential and leads to up or downregulation of reactive oxygen species (ROS), adenosine triphosphate (ATP) (9), (3) and calcium (Ca^2+^) (1). These molecules trigger the activation of transcription factors (e.g., NF-*κβ*, Nrf2 and activator protein-1[AP-1]) (10), changes in protein expression and release of cytokines and growth factors (11). The exact effects that follows are hard to predict: it includes altered mitochondrial activity (12), gene expression (1, 13, 14), promotion of anti-inflammatory response (3) and cell proliferation (15). ROS, for example, leads to apoptosis, if found in great amounts, and may also increase proliferation at lower levels. Therefore, investigating the modulation of these molecules activity by PBM and its connection with changes in metabolism and physiological effects, within the same conditions of illumination and cell type, is fundamental.

Glucose is the primary fuel of cellular respiration; its catabolism reduces the electron carriers by transferring electrons to FAD molecules, producing FADH_2_ and NAD coenzymes, providing NADH (16). The NADH and FADH_2_ are oxidized, respectively, to NAD^+^ and FAD at complexes I and II of the electron transport chain, producing an electrical potential that results in a donation of electrons to molecular oxygen and phosphorylation of adenosine diphosphate (ADP) by the ATP synthase enzyme (17). Generally, lower oxygen concentrations shift the glucose catabolism to anaerobic glycolysis, which converts glucose to lactate instead of pyruvate, supplying enough energy for the maintenance of cellular processes (18). The glycolytic pathway takes place at the cytosol resulting in ATP generation and oxidation of phosphoenolpyruvate to pyruvate.

In non-cancer cells, this pathway can either provide enough energy to cells under hypoxic conditions or supply the citric acid cycle with pyruvate to produce mitochondrial ATP by oxidative phosphorylation (OXPHOS) (19). The formed NADH and FAD of these coenzymes present an intrinsic fluorescence, which allows the redox ratio (RR) of the cell to be calculated optically by FAD/ [NADH + FAD] fluorescence intensities (17, 20, 21). The optical redox ratio (ORR) is proportional to the balance of oxidative phosphorylation/glycolysis and can be used to monitor living tissues and cells (Figure 1) (22). Several conditions change cellular metabolism and alter this balance, such as hypoxia, high carbon demands, increased proliferation rate, and fatty acid synthesis (21). The ORR is also used to investigate cancer mechanisms since different types of tumors, and cancer cells favor glycolysis over OXPHOS, even in the presence of oxygen, a phenomenon called “Warburg effect” or aerobic glycolysis (23). Choosing aerobic glycolysis could benefit cancer cells by supplying ATP faster than oxidative phosphorylation (24) and by going through an energetic pathway that produces lower concentrations of ROS (17). It must be stated that cancer cells can favor oxidative metabolism over aerobic glycolysis, for reasons not fully elucidated. Highly invasive tumor cells, for example, have shown modulation of the glucose metabolic pathway depending on the site of metastasis (25–27). Oral cancer is one of them, and its location is convenient to make optical measurements and an ORR analysis. Previous studies have shown that it is possible to differentiate healthy tissue, hyperplasia, and dysplasia with this technique *in vivo*, which shows its potential to monitor metabolism changes in the tumor (28).

**Fig.1.**
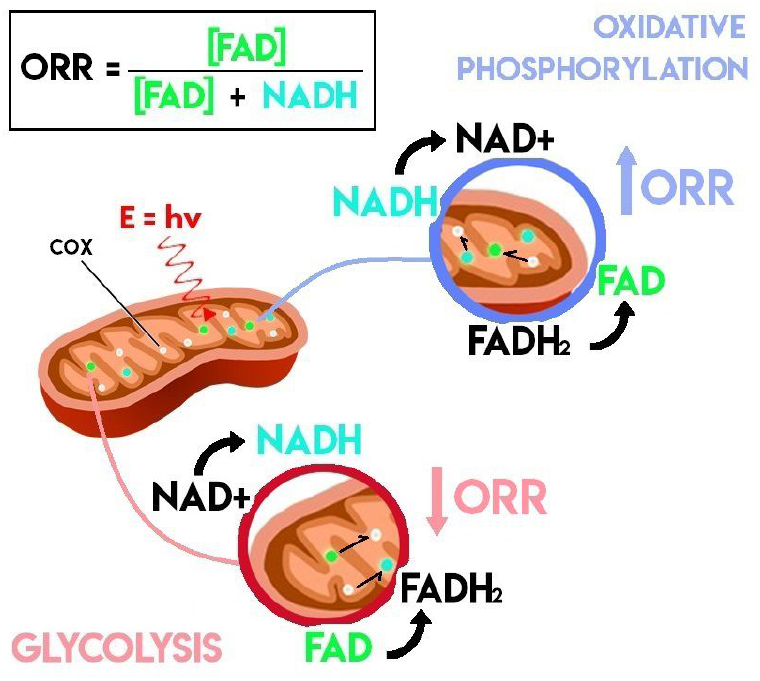
Schematics of the optical redox ratio (ORR) in mitochondria, following light absorption by cytochrome c oxidase (COX), and its correlation to the ATP generation pathway. Normal cells produce ATP by oxidative phosphorylation (OXPHOS) in normoxic conditions. Within the mitochondria, NADH and FADH_2_ are oxidized, respectively, to NAD^+^ and FAD, increasing RR. In hypoxic conditions, cells use the glycolysis pathway to supply ATP. Glycolysis reduces the electron carriers NAD^+^ and FAD to NADH and FADH_2_, respectively, lowering the ORR.

Therefore, in order to increase PBM acceptance, it is fundamental to investigate its effects on the metabolism of cancer cells, since it is a modality clinically used to treat and prevent side effects, such as mucositis, in cancer patients undergoing radio and chemotherapy. Despite studies on the activation of a few pathways and the regulation of important molecules alone do exist, the overall PBM effect on metabolism or the existing correlations among them have not been clearly identified or understood (21). Thus, the aims of this study were to explore PBM effects on ORR and its correlation with the cell cycle, ATP levels, and ROS production, and to elucidate PBM effects related to the activation of biochemical carriers and the overall impact on the metabolism of a healthy (human dermal fibroblasts neonatal - HDFn) and cancer cell (squamous carcinoma - SCC25) lineage.

## Results

### Optical Redox Ratio Imaging

For imaging, two-photons excitation fluorescence (TPEF) microscopy allowed the acquisition of high resolution images of depth sectioning without the need for a confocal pinhole, since TPEF is a non-linear light process limited to the focal plane, which also spares any damage to surrounding tissue or cells (21, 29). Figure 2 shows the NADH (blue) and FAD (green) fluorescence by TPEF microscopy and the merged image (red) indicating the ORR of the SCC-25 cells (Figure 2a-c) and HDFn (Figure 2d-f).

**Fig.2.**
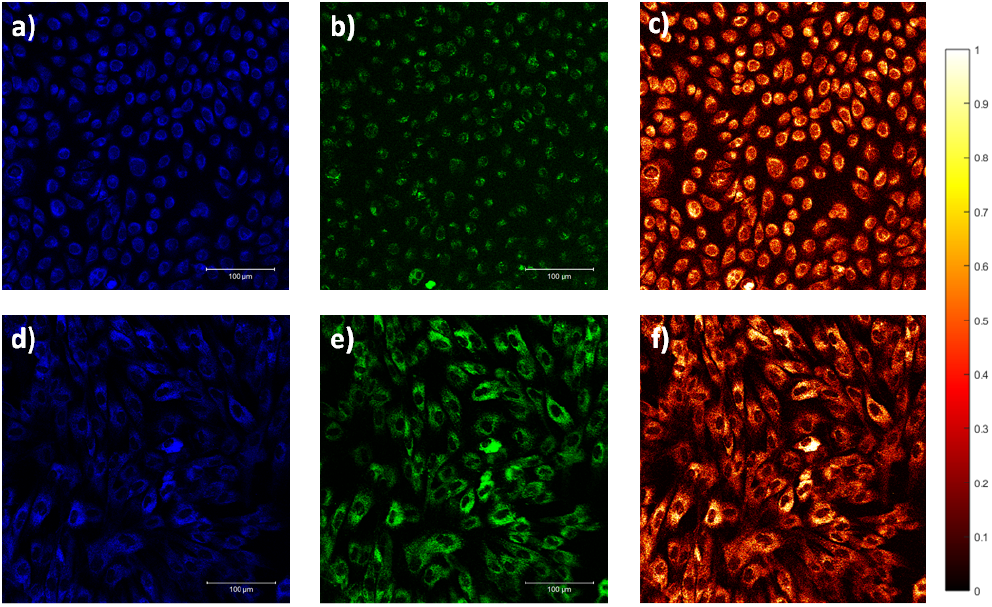
Fluorescence microscopy of SCC-25 (a-c) and HDFn (d-f) cells. The false color blue images (a and d) correspond to NADH fluorescence, false color green (b and e) correspond to FAD fluorescence and the false color red (c and f) are the calculated optical redox ratio image.

From the results shown in Figure 3a, it is evident that fibroblasts present a higher ORR than carcinoma cells. This is rational since normal cells favor oxidative phosphorylation (↟FAD/(↟FAD+↓NADH)) over glycolysis (↓FAD/(↓FAD+↟NADH)) and is consistent with previous observations (17, 21). Regarding PBM, illumination did not show a significant effect on HFDn ORR value, however, it decreased the ratio of SCC-25 cells by 10%, indicating increased glucose catabolism. Additionally, cell-to-cell ORR variability was calculated using a region of interest (ROI) mask to compute the mean redox ratio of a single cell. It is noticeable that the variability shown is greater for SCC-25 cells than for healthy HDFn cells. This is consistent with the fact that some tumor cells, presenting a more metastatic potential, contradict the Warburg effect, (17) which consists in the preferential metabolism of glucose to lactate, independent of oxygen presence, by cancer cells (30). Another interesting observation is that PBM reduced the variability in both cell lines, despite not causing a difference in the ORR mean of HDFn cells. This means that the balance of oxidative phosphorylation/glycolysis among the population became more homogeneous after illumination. If we combine this result with the decrease in the mean of SCC-25 ORR, it is possible to raise the hypothesis that PBM induces a shift towards glucose catabolism in cells that previously presented a higher rate of OXPHOS. For HDFn cells, the decrease in variability could be related to PBM producing slightly different effects according to the state of a cell, upregulating OXPHOS in cells presenting a lower redox state and decreasing glycolysis in the ones that favored it instead of OXPHOS.

**Fig.3.**
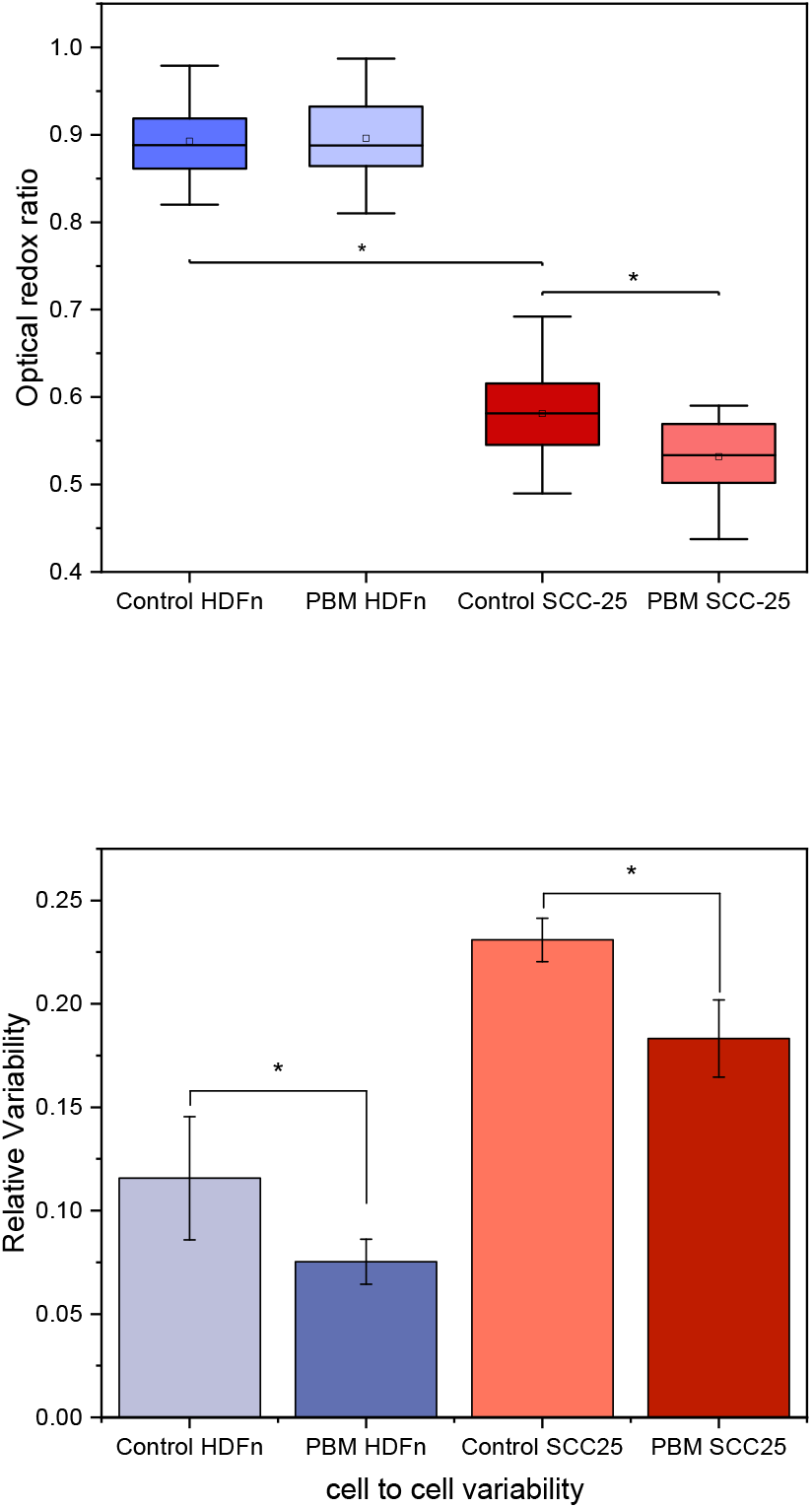
(a) Mean optical redox ratio of HDFn and SCC-25 cells, control and PBM groups. (b) Cell-to-cell relative variability in the redox ratio for SCC-25 and HDFn cells, control and PBM. *p < 0.05

### Glycolysis

Glycolysis results after 4 hr of PBM are shown in Figure 4. It is seen in Figure 4a that fibroblasts present a lower baseline for glycolysis than the tumor cell line, as expected due to the Warburg effect observed in cancer cells. The PBM caused an increase in this parameter in both cell lines (Fig 4b and 4c), in a similar proportion. As HDFn cells did not present a difference in ORR after PBM we conclude that OXPHOS increased as well, and the balance was not altered. The SCC-25 cells showed a decrease in ORR and an increase in glycolysis, making it possible to infer that OXPHOS was not affected, or had a slight decrease.

**Fig.4.**
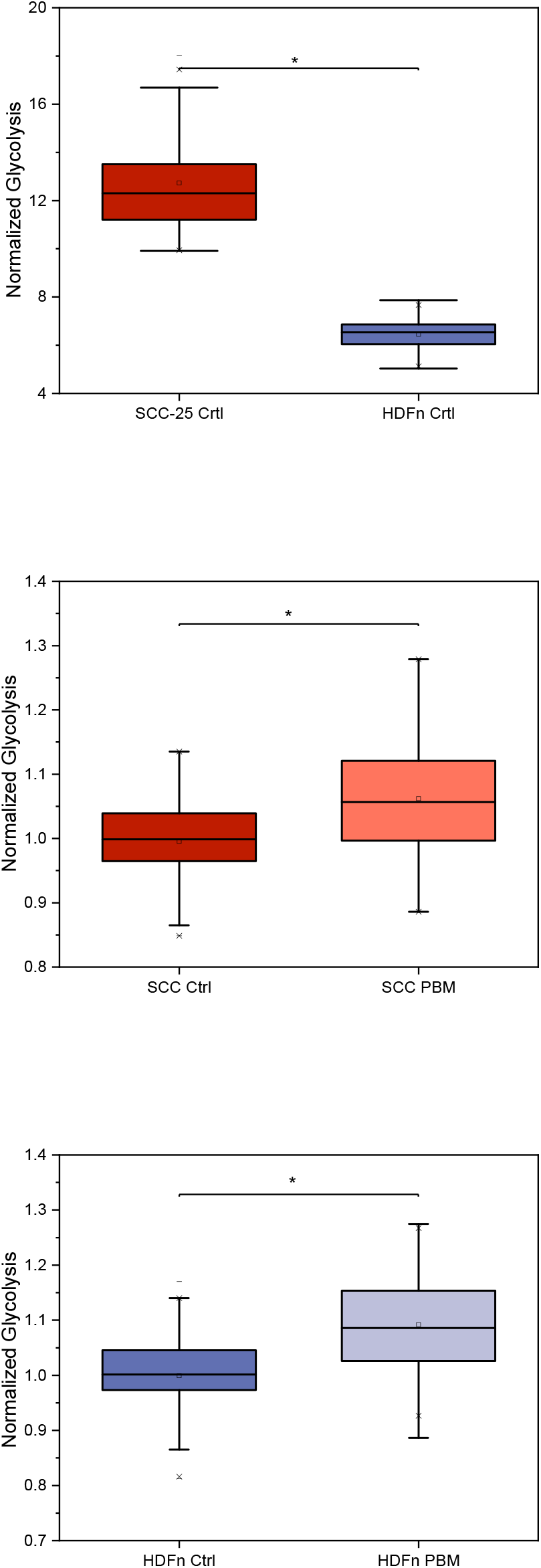
Glycolysis experiment assay. (a) Baseline of glycolysis for SCC-25 and HDFn cell line showing that the tumor cell line (SCC-25) has a greater baseline for glycolysis when compared to HDFn cells. This result was expected and could be related to the Warburg effect. (b) Glycolysis quantification after PBM for SCC-25 cell line and (c) for HDFn cell line. Results show that PBM did not influence the glycolysis rate of normal cells but increased the rate of tumor cells. * *p* < 0.05

### Metabolic activity assessment by 3-(4,5-dimethylthiazol-2-yl)-2,5-diphenyltetrazolium bromide (MTT) assay

Cell viability was assessed by the MTT assay 4 h and 24 h after PBM and is shown in Figure 5. Since this assay is used to measure cell viability based on cell metabolism, cell counting was performed to confirm the results from MTT and showed a good correlation (see SI). It is possible to observe a difference in cell viability 4 h after PBM in both cell lines alongside similar cell counting, which indicates a change in metabolism in both cells. Mitochondrial activity was increased in fibroblasts (Fig. 5a) and decreased in SCC-25 cells (Fig. 5b). At 24 h, it was observed that PBM induced proliferation in fibroblasts, as both MTT and cell counting increased. However, there was no significant change in the tumor population.

**Fig. 5.**
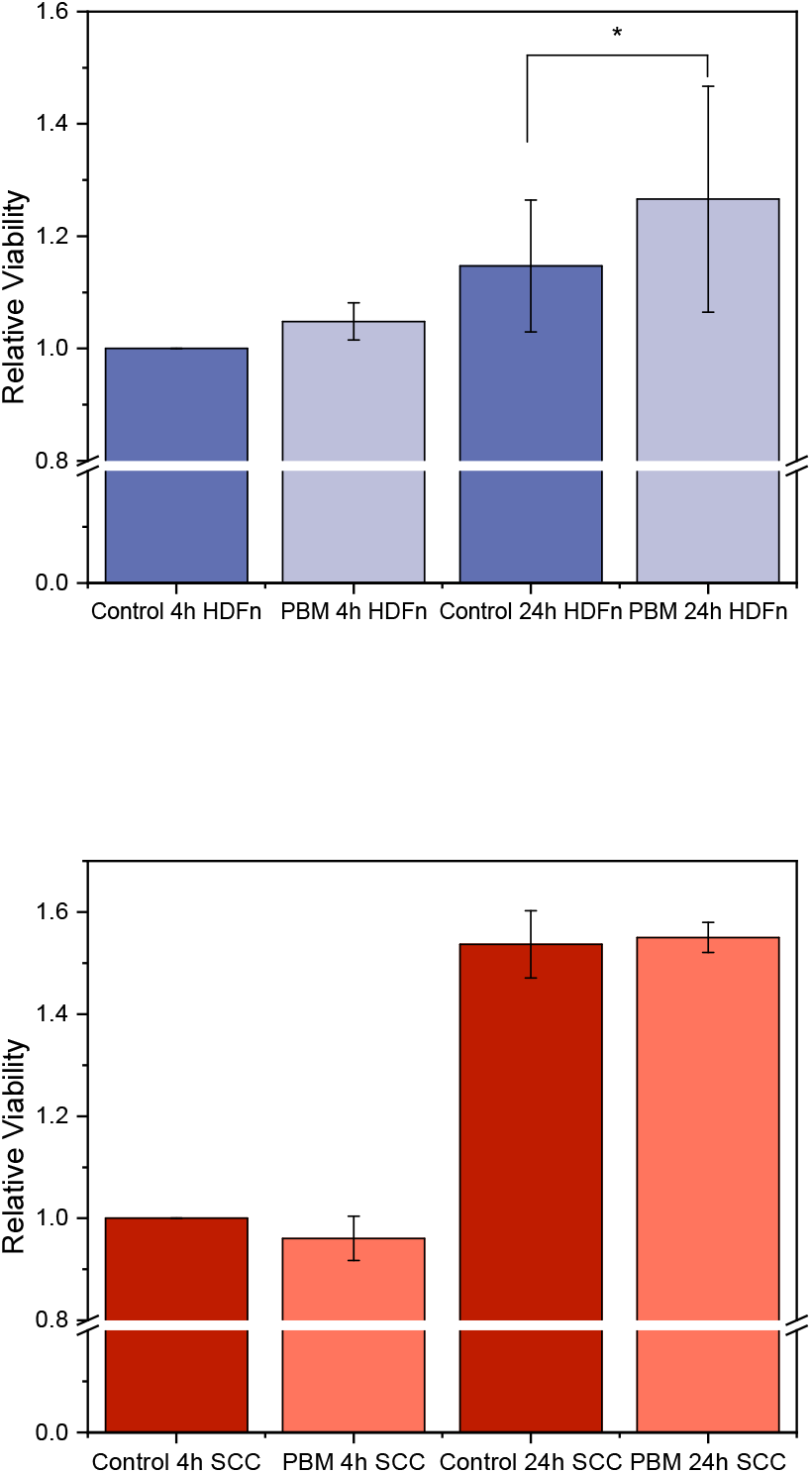
Metabolic activity by MTT assay 4 h and 24 h after PBM. (a) shows the HDFn cell line viability of control samples and illuminated samples, indicating that PBM induced cell proliferation in fibroblasts after 24 h. (b) Cell viability assay for SCC-25 cells. When compared to control, no cell proliferation was observed in tumor line 24 h after illumination.

### ROS and ATP Assay

The ROS quantification after PBM was performed by flow cytometry to investigate if its production correlated to illumination (Figure 6). Figure 6a shows the ratio of mean intensities between PBM and the control of each cell line. In fibroblasts, no significant (p > 0.05) changes were found among the samples. In SCC-25, however, a statistically significant (p < 0.05) increase of about 30% was observed after PBM. This suggests that ROS could play an important role in mediating PBM effects in tumor cells but not in normal fibroblasts. One common consequence of several pathways initiated by ROS is increased ATP production. As seen in Figure 6b, endogenous ATP increased within the 24 h-after PBM period evaluated for SCC-25 and HDFn cells, even though kinetics differed among the cell lines. The SCC-25 cells presented a peak of 1.25 units compared to the control at 4 h after PBM while fibroblasts modestly increased ATP by 7 % 6 h after PBM. Interestingly, both cells showed a decrease immediately after its ATP peaks, indicating consumption by energy demanding processes.

**Fig. 6.**
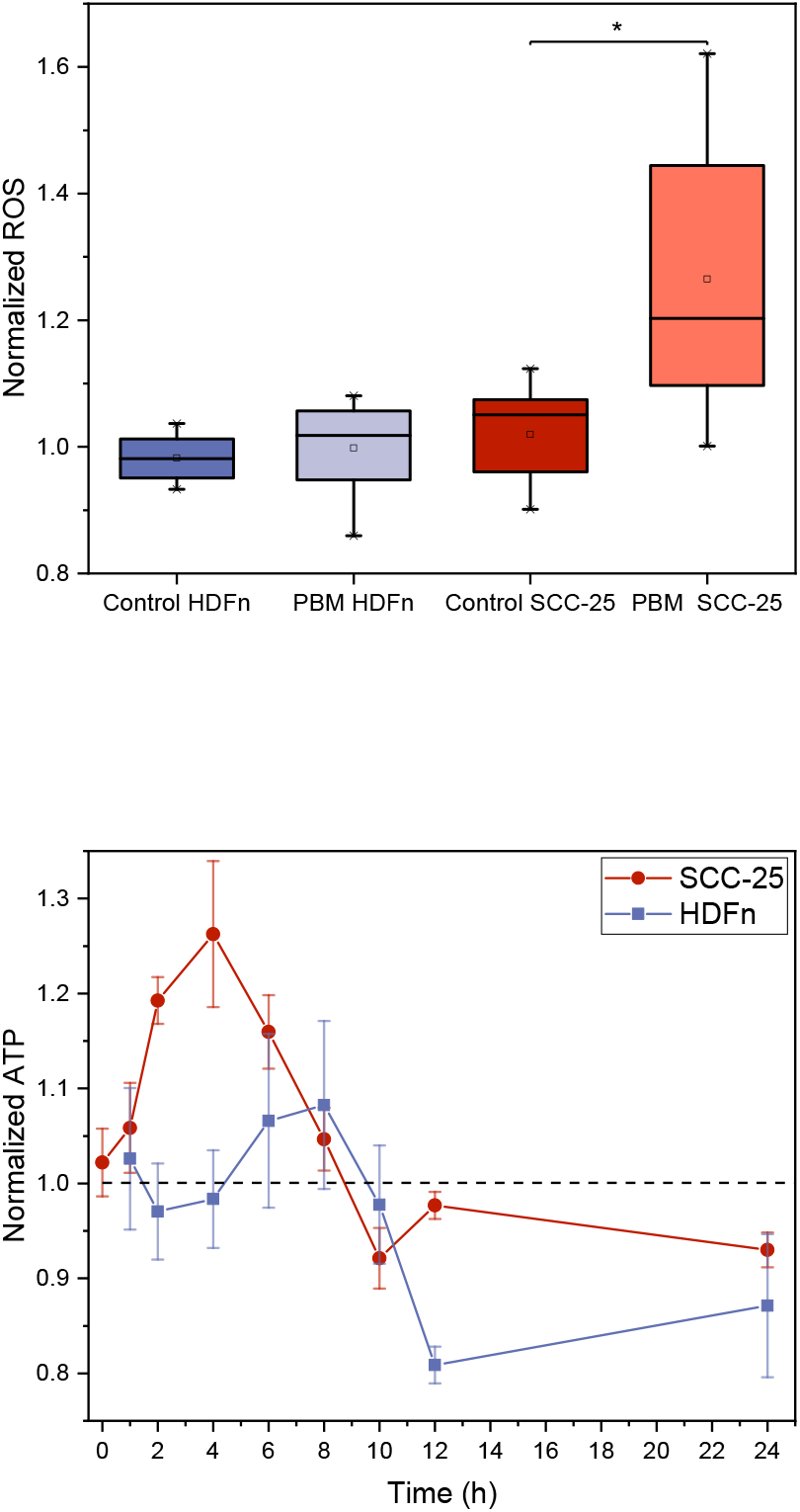
ROS and ATP assay, (a) The ROS production assay indicating that PBM induced ROS production in the SCC-25 cell line (p < 0.05), but not in fibroblasts. (b) The ATP production of both cell lines, indicating that SCC-25 (red-dot) cells increased a peak of 1.25 units 4 h after PBM, as fibroblasts modestly increased 7 % after 6 h (grey-square).

### Cell cycle assessment

Cell cycle was evaluated by flow cytometry 8 h and 24 h after PBM or sham treatment. The proportion of cells in G2/M after PBM relative to the control is shown in Figure 7. Cells in G0/G1 and S phase were not statistically different. It was observed that both fibroblast and tumor cells increased mitosis in a linear manner and at the same rate, reaching a 20 % increase in 24 h.

**Fig. 7.**
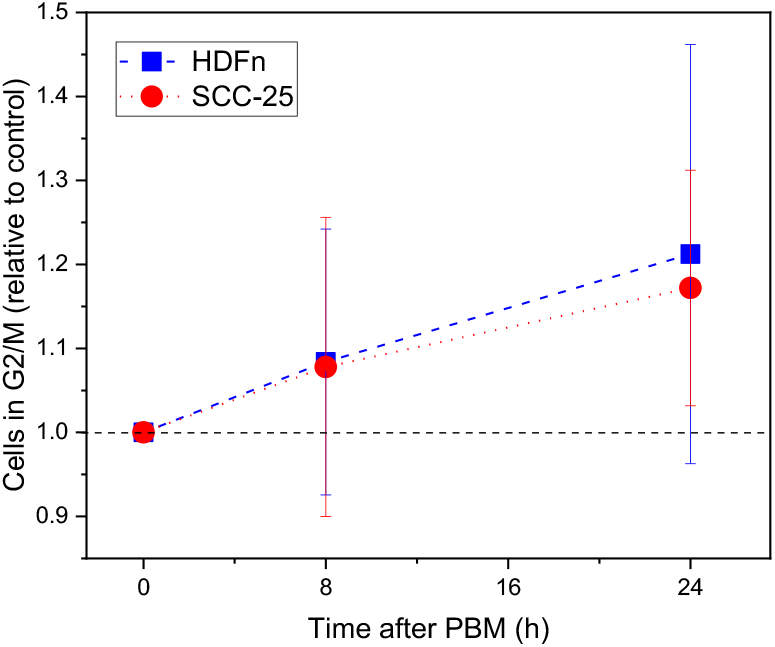
Cell cycle assessment by flow cytometry Illuminated samples of HDFn (blue-square line) and SCC-25 (red-dot line) linearly increased the mitosis rate up to 20% after 24 h when compared to controls.

## A. Discussion

Photobiomodulation is the use of light, mainly in the red and near-infrared regions, for a variety of purposes. It is promising since it is a non-invasive and an affordable technique already used to reduce inflammatory conditions (3), in the treatment of arthritis (31) and wound healing (32), among others, resulting in pain relief and modulation of expression of genes related to the inflammatory response (6, 33, 34). The PBM encompasses such a broad spectrum of illumination protocols, parameters, and uses; its mechanism of action is not fully understood. This causes skepticism from the medical community and limits its impact. As stated by Stephen Sonis (35), until we obtain enough data we cannot answer whether we should avoid PBM in head and neck cancer tumors or not. So it is fundamental to understand these effects to ensure the safety of this technique and explore its potential in enhancing cancer treatment.

In this study, we investigated the effects of PBM on the metabolism of healthy (HDFn) and cancer cells (SCC-25) *in vitro* and revealed that their pathways are different. It was also established that ORR evaluation by TPEF is a technique that is sensitive enough to significantly detect slight changes caused by PBM. Thus, it is a powerful tool to investigate metabolism modulation in both cancer and normal cells. The PBM illumination protocol was based on previous mucositis studies (36–38) and the results are summarized in Figure 8. In fibroblasts cells, no changes in the redox state were observed 4 h after illumination despite increased glycolysis displayed by a different method. Therefore, both forms of respiration must have increased at the same rate in these cells, maintaining the ratio constant. In SCC-25 cells, a lower ORR shows that PBM modulates a shift in the redox state of the cells towards glycolysis.

**Fig. 8.**
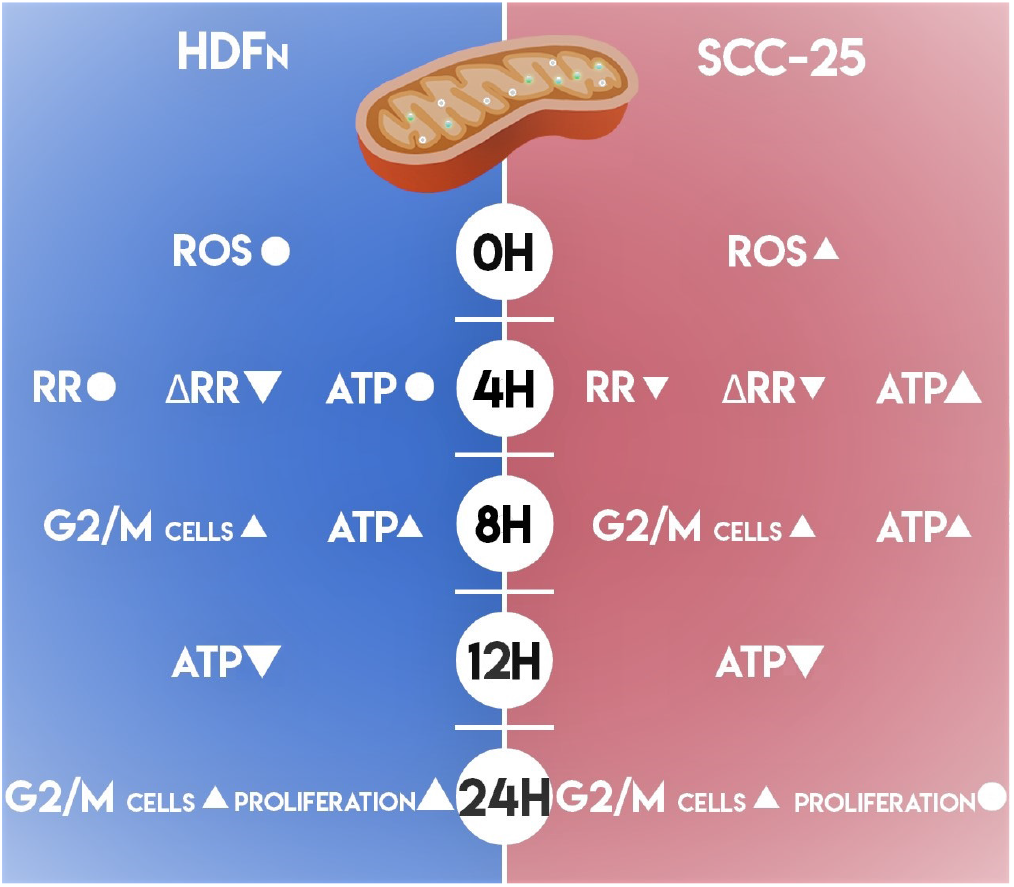
Summary of HDFn and SCC-25 modulations caused by PBM indicating increase, decrease or no change in reactive oxygen species (ROS), redox ratio (RR), redox ratio variability (ΔRR), adenosine triphosphate (ATP), number of cells in G2/M and proliferation, compared to its respective controls.

Previous work by Heymann and colleagues reported a PBM-induced decrease in the redox ratio, measured by the extra-cellular flux assay, along with increased proliferation in HeLa cells, using 670 nm and 12 J/cm^2^ (39). Since the illumination protocol and cell type were different, but the effect similar, it might be a common effect of PBM in tumors. Its consequences in cancer cells need to be investigated since it may correlate to the Warburg effect and its therapeutic implications, such as tumor aggressiveness shown by Li et. al. (40). Additional evidence that arose from the ORR analysis is that PBM may have different effects and mechanism of action depending on the previous redox state of the cells. This was shown by decreased variability in ORR values 4 h after illumination, in both cell lines, caused by a decrease in the highest and an increase in the lowest values of ORR. It suggests that PBM acts differently according to the cells. In this instance, it may be more effective to the ones that differ from the mean redox state of the population.

Beyond the differences in ORR, fibroblasts and SCC-25 cells, these seem to have distinguished pathways that initiate the cascade of events that characterize PBM. ROS is known to be an important biomarker that induces apoptosis if found in high concentrations, and modulates pro-survival and proliferation effects at low concentrations (41). In this study, ROS concentration increased only in SCC-25 cells, indicating that PBM acts by a different pathway in fibroblasts. Engel and colleagues showed increased catalase in fibroblasts after PBM scavenged ROS. Therefore, they suggested that lineage-specific differences maintain homeostatic redox status within each cell type (41).

Nevertheless, ATP levels were increased in both cell lines after PBM. Chen et. al. showed in fibroblasts, that ATP increase after PBM is not altered with the addition of antioxidants. Despite showing an increase in ROS that was not seen in our study, both results suggest that ATP synthesis after PBM is not dependent on ROS signaling (42). ATP kinetics after PBM, however, have not been investigated yet. Such an investigation is important because end-point measurements can lead to false conclusions. For example, ATP increase in fibroblasts was only seen 8 h after PBM while its peak for SCC-25 cells was seen at 4 h. At 12 h, ATP levels were lower when compared to the non-illuminated groups for both cells lines. This indicates higher ATP demands from processes induced by PBM, such as protein synthesis and DNA replication involved in proliferation, or a mechanism of feedback that tends to suppress the effects caused by light.

In fact, we observed an increase in G2/M fraction for both cells at 8 and 12 h after PBM. However, it did not result in increased proliferation in the tumor cell line but did in HDFn cells. This is an encouraging result that supports the evidence that PBM does not affect tumor growth (43). Schartinger et al. reported similar results using 660 nm, an increase in fibroblasts but a decrease in SCC-25 cells (44). This indicates regarding proliferation, that PBM effects are similar for multiple wavelengths. Regarding cell cycle, they observed an increase fraction in cell cycle G1 and S phases, but did not report the time after PBM in which the measurement was performed. In contrast, Sperandio et al. observed increased proliferation in SCC-25 cells for both 660 and 780 nm, at 24 h (780 nm, 6.15 *J/cm*^2^) and 48 h (660 and 780 nm, 3.07 *J/cm^2^*) after illumination (45). Certainly, further studies need to be conducted to understand if PBM stimulates proliferation in tumors, and under what conditions, in order to advance the reliability and security of its applications in cancer.

Therefore, it was demonstrated that PBM with 5 *J/cm*^2^ at 780 nm alters the metabolism of fibroblasts and HNSCC cells, but in different pathways and kinetics. Its mechanism of action needs to be further investigated to improve the understanding of these differences. For that, studies in more complex models, 3D cell cultures and *in vivo*, need to be conducted. So the influence of the extracellular matrix, spatial fluence distribution, surrounding tissues, immune and vascular response, among others, can be evaluated. Then, it may be possible to explore PBM mechanisms to improve cancer treatments, or avoid applications involving tumors to prevent negative effects. Additionally, TPEF was depicted as a powerful tool to evaluate redox state after PBM. It is a sensitive technique that allows the assessment of small redox ratio differences and variability among cells. It is also nondestructive, so the sample can be used after measurements, and it can be combined with other fluorescent markers.

## B. Material and Methods

### Cell Culture

Human dermal fibroblasts neonatal (HDFn) and squamous carcinoma SCC-25 (American Type Culture Collection - ATCC), Wesel, Germany), were cultivated at 37°C in humidified 5% CO2 atmosphere in Dulbecco’s modified Eagle medium (DMEM) and DMEM/Ham’s (Cultilab), respectively. Media were supplemented with 10% (v/v) Fetal Bovine Serum (Cultilab, Brazil) and to DMEM/Ham’s hydrocortisone was added (Sigma-Aldrich, USA) at a concentration of 400 ng/ml.

### Illumination protocol

PBM groups were illuminated using a custom-made LED array device emitting at 780 nm with an irradiance of 30 *mW/cm^2^* and a total fluence rate of 5 *J/cm^2^.* (46) The control groups were sham treated.

### Optical Redox Ratio Imaging

Cells were plated on a 35 mm glass bottom dish (Greiner Bio-One, Germany) at a density of 5 x 10^5^ cells and let in a heated chamber (37°C, 5%, CO_2_) overnight. Four hours after PBM, cells were washed twice in PBS and the images were performed on an inverted fluorescence confocal microscope (Zeiss - LSM780, Zeiss, Germany) equipped with a Ti:Sapphire tunable laser source (Chameleon Vision II, Coherent Inc., USA). The laser excitation source was tuned to 755 nm (NADH excitation) or 860 nm (FAD excitation), and images were acquired in the channel mode of the microscope with 440 - 480 nm (NADH fluorescence) or 500 - 550 nm (FAD fluorescence) wavelength range, respectively. Images (1024 x 1024 pixels; 8-bit depth; 425 *μm* x 425 *μm*) were acquired using a 20x objective (NA = 0.8). For each condition, two plates were prepared and 10 fields were imaged for each one. Two independent experiments were performed, resulting in N=40. To calculate cell-to-cell ORR variability, a region of interest (ROI) was selected and used to create a mask to compute the mean ORR of a single cell. The mask was created manually from the FAD image. Three cells of each field were analyzed, resulting in the analysis of 120 cells per group. To ensure that the same dish would yield the same result, two dishes were calculated twice, using different cells from the field. Then, the ‘variability’, defined as the relative standard deviation (standard deviation/mean ORR), was calculated for each dish. Therefore, the error of this parameter is the standard deviation of its values for four dishes. All images were acquired using Zen 2010 software (Zeiss, Germany). A control plate was imaged every in experiment-day in order to normalize some microscope variations. Image analysis was performed using MATLAB (MathWorks, USA) and the redox images were created by computing pixel-wise ratios of FAD/(NADH + FAD) fluorescence. For statistical analysis and bar plot presentation, the average redox ratios of cell plates were calculated by separately computing the average FAD and NADH intensities from the respective images and taking the ratio of these values.

### Glycolysis assay

Glycolysis was assessed with a fluorescent kit (Abcam ab197244, Abcam, USA) following manufacturer protocols. 2×10^4^ cells/well were seeded in 96-well opaque black walls 24 h prior illumination, in 6 replicates per group. Then, 1h after PBM, CO_2_ was removed from the incubator and at 4h after PBM wells were washed twice with Respiration Buffer and 15 *μl* of Glycolysis assay reagent in 100 ul of Buffer was added to each well. Fluorescence (ex/em: 380/615 nm) was measured with a SpectraMax M5 Multi-Mode Microplate Reader (Molecular Devices, USA) for 2h in 1.5 min intervals. The means correspond to two independent experiments.

### Metabolic activity assessment by MTT assay

Metabolic activity was assessed at 4h and 24h after PBM. Cells were seeded in triplicate for each condition in 24-wells plates at a density of 1×10^5^ per well (500 *μl*) and illuminated the following day according to the parameters mentioned above. After 4h or 24h, medium was replaced by 250 *μl* of new media with 3-(4,5-dimethylthiazol-2-yl)-2,5-diphenyltetrazolium bromide (MTT) (5 *μ*g/ml) and incubated for 3h, until 1 ml of DMSO was added and absorbance was measured at 570 nm in a microplate reader (Multiskan™ FC Microplate Photometer – ThermoFisher Scientific, USA). Each experiment was performed three independent times. To confirm whether the results from MTT resulted proliferation and viability, a trypan blue exclusion assay was performed in quadruplicate, in the same conditions.

### ROS Assay

Quantification of ROS after PBM was performed by flow cytometry assessment using DCFH-DA. For the assay, a 1×10^6^ cells per ml suspension was made in phenol and FBS free medium. Triplicates of 250 *μl* of the cell suspension were illuminated in a 24-wells plate with a dose of 5 J/cm^2^ at 780 nm in a black 24-wells plate with clear bottom. Samples were immediately incubated with 250 *μl* of DCFDA solution, resulting in a concentration of 25 *μM*, for 30 minutes at room temperature in the dark and assessed by flow cytometry (BD, C6 Accuri Plus, USA) at an excitation/emission of 492–495 nm/517–527 nm.

### ATP Assay

Cells were seeded in 96-well plates at a 2×10^4^ cells/well density and incubated at 37°C, 5% CO_2_ for 24h prior the ATP assay, performed with the ATP bioluminescent assay kit (Sigma–Aldrich, USA). Plates were illuminated or sham-illuminated and at a specific time after that ranged from 1-24h supernatant was removed, wells were washed twice with PBS and 100 *μl* of Releasing Reagent were added. The working solution was prepared as indicated (10% of ATP Mix Working Solution in ATP Mix Dilution Buffer). Immediately prior to the bioluminescent reading, 100 *μl* was added to the wells with a multi-channel pipette to ensure all wells were incubated simultaneously and only 6 wells were read at a time. The luminescence was measured with a SpectraMax M5 Multi-Mode Microplate Reader (Molecular Devices, USA). Experiments were repeated three times with 6 replicates per group (N=18).

### Cell cycle assessment

Cell cycle evaluation was performed by flow cytometry analysis using propidium iodide (PI). PBM was performed in 24-wells plate as described previously, in triplicate. Then, at 0h, 8h and 24h after illumination, cells were collected and fixed in ice-cold 70 % ethanol at −20°C for at least 24 h, then washed with PBS and stained with PI (50 *μg* PI/ml in PBS, BD Biosciences) containing 0.1 mg/ml RNase (Sigma–Aldrich, USA) for 40 min. Samples were analyzed in an Accuri C6 flow cytometer (BD Biosciences, USA) in triplicate and cell cycle was determined using FlowJo software univariate analysis (BD Biosciences, USA). Two independent experiments were performed, with a final N=12.

### Statistical analysis

The data were plotted using boxplot with a whisker of 1-99 or represented as means ± standard deviation and were analyzed using the commercially available software Origin 2018 (Origin Lab., USA). One-way analysis of variance (ANOVA) was used among the categories “HDFn” and “SCC-25” cells and “Control” and “PBM” for the ORR measurements. For experiments that we compared only “PBM” and “Control” independently for the same cell line, a single ANOVA test was performed. Differences were considered as statistically significant at p<0.05. Asterisks placed above bars indicate statistical significance.

## ACKNOWLEDGEMENTS

The authors thank Camila Bramorski and André Longo for the support with the figures and acknowledge the support provided by Brazilian Funding Agencies: Coordenação de Aperfeiçoamento de Pessoal de Nível Superior - Brasil (CAPES) - Finance Code 001; CNPq (465360/2014-9) and São Paulo Research Foundation (FAPESP) grants: 2009/54035-4 (EMU); 2013/07276-1 (CePOF); 2014/50857-8 (INCT); 2017/14182-4 (CMGF scholarship).

